# Representation Learning of Resting State fMRI with Variational Autoencoder

**DOI:** 10.1101/2020.06.16.155937

**Authors:** Jung-Hoon Kim, Yizhen Zhang, Kuan Han, Zheyu Wen, Minkyu Choi, Zhongming Liu

## Abstract

Resting state functional magnetic resonance imaging (rsfMRI) data exhibits complex but structured patterns. However, the underlying origins are unclear and entangled in rsfMRI data. Here we establish a variational auto-encoder, as a generative model trainable with unsupervised learning, to disentangle the unknown sources of rsfMRI activity. After being trained with large data from the Human Connectome Project, the model has learned to represent and generate patterns of cortical activity and connectivity using latent variables. The latent representation and its trajectory represent the spatiotemporal characteristics of rsfMRI activity. The latent variables reflect the principal gradients of the latent trajectory and drive activity changes in cortical networks. Latent representations are clustered by both individuals and brain states. Representational geometry captured as covariance or correlation between latent variables, rather than cortical connectivity, can be used as a more reliable feature to accurately identify subjects from a large group, even if only a short period of data is available per subjects.

## INTRODUCTION

The brain is active even at rest, showing complex activity patterns measurable with resting state fMRI (rsfMRI) (Fox and Raichle, 2007). It is widely recognized that rsfMRI activity is shaped by how the brain is wired, or the brain connectome (Sporns et al., 2005). Inter-regional correlations of rsfMRI activity are often used to report functional connectivity (Biswal et al., 1995) and map brain networks for individuals (Finn et al., 2015) or populations in various behavioral (Smith et al., 2009) or disease states (Fox et al., 2014). However, it remains largely unclear where rsfMRI activity comes from (Leopold and Maier, 2012; Lu et al., 2019), whereas understanding its origins is critical to interpretation of any rsfMRI pattern or dynamics (Winder et al., 2017).

Prior findings suggest a multitude of sources (or causes) for rsfMRI activity (Bianciardi et al., 2009), including but not limited to fluctuations in neurophysiology (Mantini et al., 2007), arousal (Chang et al., 2016), unconstrained cognition (Chou et al., 2017), non-neuronal physiology (Birn et al., 2008), head motion (Power et al., 2014) etc. These sources only partially account for rsfMRI activity and may be entangled not only among themselves but also with other sources that are left out simply because they are hard to specify or probe in a task-free state (Leopold and Maier, 2012). An inclusive study would benefit from using a data-driven approach to uncover and disentangle all plausible but hidden sources from rsfMRI data itself, without having to presume the sources to whatever are experimentally observable. To be effective, such an approach should be able to infer sources from rsfMRI data and generate new rsfMRI data from sources, while being able to account for complex and nonlinear relationships between the sources and the data.

These requirements lead us to deep learning, or representation learning with deep neural networks (LeCun et al., 2015), as a nonlinear method for blind source separation, in contrast to its linear counterparts, e.g. independent component analysis (Beckmann and Smith, 2004; Calhoun et al., 2001; Smith et al., 2012). For brain research, deep learning models has provided testable models of the brain in terms of neural computation for sensory and language processing (Han et al., 2019; Kell et al., 2018; Khaligh-Razavi and Kriegeskorte, 2014; Richards et al., 2019; Wen et al., 2018; Yamins and DiCarlo, 2016; Zhang et al., 2020). Deep learning has also been increasingly used as a generic family of machine learning tools to learn features from fMRI data. See (Khosla et al., 2019b) for a review. Most applications are in the regime of supervised learning. Typically, a neural network takes an fMRI-based input data and is trained to generate an output that optimally matches the ground truth for a task, such as individual identification (Chen and Hu, 2018; Wang et al., 2019), prediction of gender, age, or intelligence (Fan et al., 2020; Gadgil et al., 2020; Plis et al., 2014), disease classification (Seo et al., 2019; Suk et al., 2016; Wang et al., 2020; Yang et al., 2019; Zou et al., 2017). The labels required for supervised learning are often orders of magnitude smaller in size than the fMRI data itself, which has a high dimension in both space and time. As a result, the prior studies often limit the model capacity by using a shallow network and/or limit the input data to activity at the region of interest (ROI) level (Chen and Hu, 2018; Dvornek et al., 2018; Koppe et al., 2019; Matsubara et al., 2019; Suk et al., 2016; Wang et al., 2019; Wang et al., 2020) or reduce it to functional connectivity (D’Souza et al., 2019; Fan et al., 2020; Kawahara et al., 2017; Kim and Lee, 2016; Riaz et al., 2020; Seo et al., 2019; Venkatesh et al., 2019; Yang et al., 2019; Zhao et al., 2018). It is also uncertain to what extent representations learned for a specific task would be generalizable to other tasks. It is further debatable whether deep neural networks with supervised learning are currently superior to more conventional and simpler methods (He et al., 2020).

For these considerations, unsupervised learning is more preferable for uncovering the underlying causes that drive intrinsic brain activity regardless of any task or disease. We choose to use the Variational Auto-Encoder (VAE) (Higgins et al., 2017; Kingma and Welling, 2013), for unsupervised learning of the increasing “big data” in rsfMRI without requiring any label or narrowly focusing on any downstream task. Unlike auto-encoder, VAE is a generative model capable of synthesizing new data similar to the training data, and it regularizes the latent space with *a priori* spherical Gaussian distributions. These properties allow the representation learned to be expressed in terms of latent variables that encode the disentangled causes of the data. Our emphasis on disentangling latent representations sets this work apart from several prior work based on the auto-encoder implemented in various forms of deep neural networks (Cui et al., 2019; Huang et al., 2017; Liu et al., 2020a; Makkie et al., 2019; Suk et al., 2016; Zhao et al., 2018). Briefly in this study, we designed and trained a VAE model to represent rsfMRI data in terms of its latent sources and tested its ability to explain and generate rsfMRI data. We characterized the time evolving trajectory of latent representation and factorized its gradients by principal components. We also visualized the representational gradients, clusters, and geometries within and across individuals, as a way to characterize brain networks and their dynamic interactions. Lastly, we tested the use of this model for characterizing individual variations and identifying individuals from their rsfMRI data (Finn et al., 2015), as a starting example of its applications.

## Methods

### Data

We used rsfMRI data from 650 healthy subjects randomly chosen from the Q2 release by HCP (Van Essen et al., 2013). For each subject, we used two sessions of rsfMRI data acquired from different days with either the right-to-left or left-to-right phase encoding. Each session included 1,200 time points separated by 0.72s. Following minimal preprocessing (Glasser et al., 2013) and automatic denoising with ICA (or the ICA-FIX) (Griffanti et al., 2014; Salimi-Khorshidi et al., 2014), we applied voxel-wise detrending (regressing out a 3^rd^-order polynomial function), bandpass filtering (from 0.01 to 0.1 Hz), and normalization (to zero mean and unitary variance). We further separated the data into three sets, including 100, 50, or 500 subjects for training, validating, or testing the VAE model, respectively. The validation dataset was used to determine the hyper-parameters used in the VAE model. The testing data were neither seen nor used by the model during training or validation. This held-out data was used to test the generalizability of the model across different datasets. For an exploratory analysis, we additionally tested the model with rsfMRI data that did not go through denoising with ICA-FIX to evaluate the model performance against presumably noisier rsfMRI data.

### Geometric reformatting

We converted the rsfMRI data from 3-D cortical surfaces to 2-D grids in order to structure the rsfMRI pattern as an image to ease the application of convolutional neural networks. As illustrated in Figure 1.a, we inflated each hemisphere to a sphere by using FreeSurfer (Fischl, 2012). For each location on the spherical surface, we used cart2sph.m in MATLAB to convert its cartesian coordinates (*x*, *y*, *z*) to spherical coordinates (*a*, *e*), which reported the azimuth and elevation angles in a range from −*π* to *π* and from 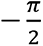 to 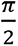, respectively. We defined a 192×192 grid to resample the spherical surface with respect to azimuth and *sin*(elevation) such that the resampled locations were uniformly distributed at approximation (Supplementary Figure 1). We used the nearest-neighbor interpolation to convert data from the 3-D surface to the 2-D grid, and vice versa.

**Figure 1.**
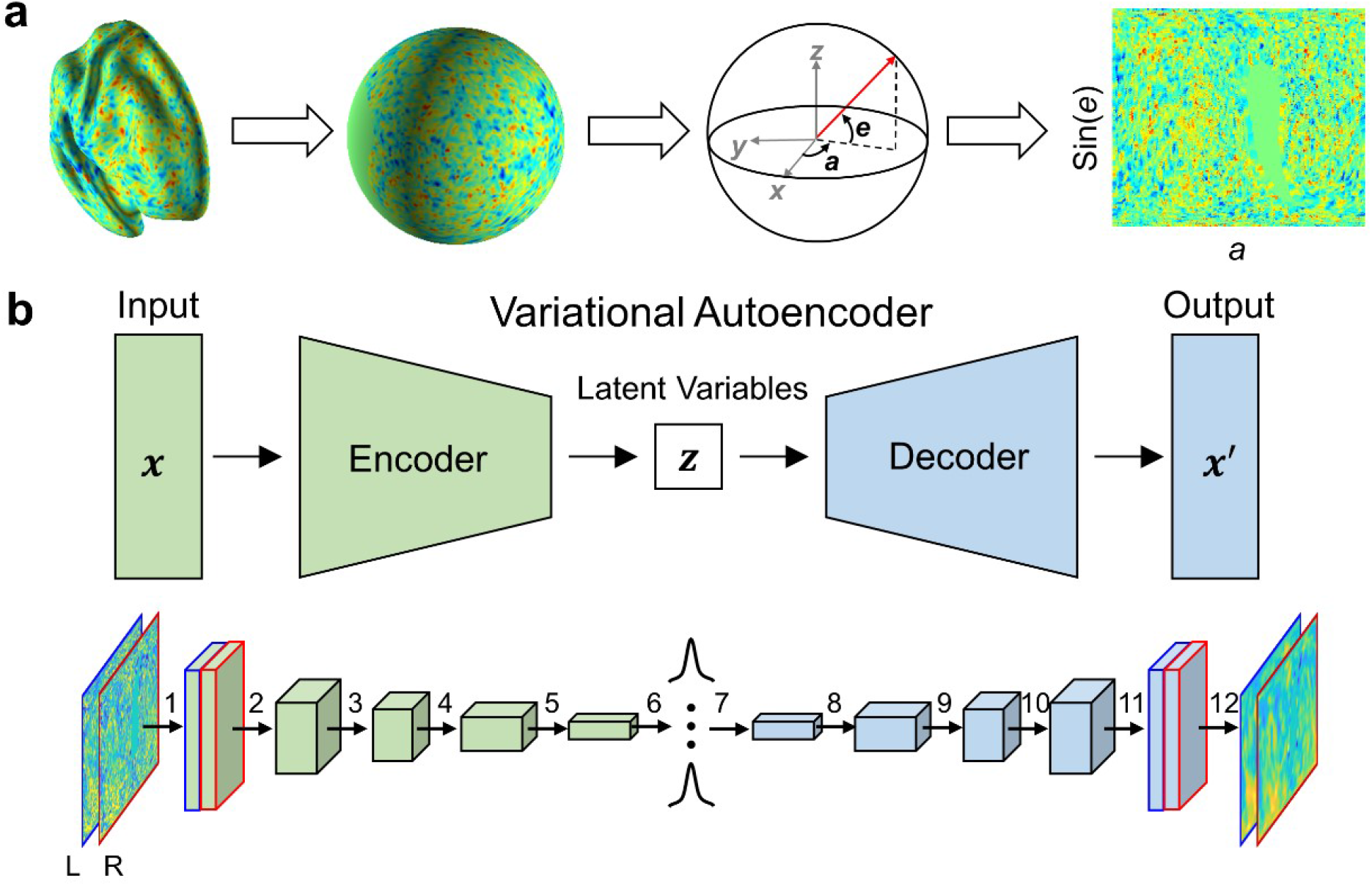
Variational Auto-Encoder (VAE). **(a) Geometric reformatting.** The cortical distribution of fMRI activity is converted onto a spherical surface and then to an image by evenly resampling the spherical surface with respect to sin(e) and a, where e and a are elevation and azimuth, respectively. **(b) Encoder-decoder architecture.** The encoder and the decoder each contain 5 convolutional layers connected in series. In the encoder, each layer (numbered from 1 to 5) outputs a feature map with the size of 96×96×64, 48×48×128, 24×24×128, 12×12×256, or 6×6×256, respectively. In layer 1, 32 kernels are applied to 192×192 flattened images of each hemisphere separately, and output feature maps are concatenated along the kernel dimension, resulting in a feature map with 64 channels. In the decoder, each layer (numbered from 8 to 12) outputs a feature map with a size of 6×6×256, 12×12×256, 24×24×128, 48×48×128, or 96×96×64, respectively. The operation at each layer is specified as follows. 1: convolution (kernel size=8, stride=2, padding=3) and rectified nonlinearity; 2-5: convolution (kernel size=4, stride=2, padding=1) and rectified nonlinearity; 6: fully connected layer and re-parameterization; 7: fully connected layer and rectified nonlinearity; 8-11: transposed convolution (kernel size=4, stride=2, padding=1) and rectified nonlinearity; 12: transposed convolution (kernel size=8, stride=2, padding=3). The blue and red boundaries highlight the input and output images for the left and right hemispheres, respectively.

### Variational autoencoder

We designed a *β*-VAE model (Higgins et al., 2017), a variation of VAE (Kingma and Welling, 2013), to learn representations of rsfMRI spatial patterns. This model included an encoder and a decoder (Figure 1.b). The encoder converted an fMRI map to a probabilistic distribution of 256 latent variables. Each latent variable was a Gaussian random variable with a mean and a standard deviation. The decoder sampled the latent distribution to reconstruct the input fMRI map or generate a new map, which appeared similar to what would be observable with fMRI. The encoder stacked five convolutional layers and one fully connected layer. Every convolutional layer applied linear convolution and rectified its output (Nair and Hinton, 2010). The 1^st^ layer applied 8×8 convolution separately to the input from each hemisphere and concatenated its output. To the feature maps concatenated across both hemispheres, the 2^nd^ through 5^th^ layers applied 4×4 convolution. Since a spherical pattern is circularly continuous with respect to the azimuth, we applied circular padding to the boundaries of azimuth for the flattened 2-D map but applied zero padding to the boundaries of elevation. Such padding was intended to avoid artifacts when otherwise applying convolution near those boundaries. The fully connected layer applied linear weighting and yielded the mean and standard deviation that described the normal distribution of each latent variable. The decoder used nearly the same architecture as the encoder but connected the layers in the reverse order for transformation from the latent space back to the input space. Figure 1.b illustrates the model architecture.

We trained the VAE model to reconstruct input while constraining the distribution of every latent variable to be close to an independent and standard normal distribution. Specifically, using the training data, we optimized the encoding parameters, *φ*, and the decoding parameters, *θ*, to minimize the loss function as below.

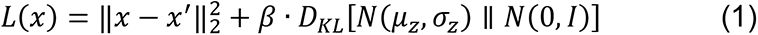

where *x* is the input data combined across the left and right hemispheres, *x*^′^ is the corresponding output from the model, *N*(*μ*_*z*_, *σ*_*z*_) is the posterior normal distribution of the latent variables, *z*, with their mean and standard deviation denoted as *μ*_*z*_ and *σ*_*z*_, *N*(0, *I*) is an independent and standard normal distribution as the prior distribution of the latent variables, *D*_*KL*_ measures the Kullback-Leibler (K-L) divergence between the posterior and prior distributions, and *β* is a hyperparameter balancing the two terms in the loss function. Part of the medial cortical surface that corresponds to corpus callosum (i.e. white matter) was excluded from training such that the learned model was intended to merely represent the activity of cortical gray matter. To train the model, we used *β* = 10 and stochastic gradient descent (batch size=128, initial learning rate=10^-4^, and 100 epochs) and Adam optimizer (Kingma and Ba, 2014) implemented in PyTorch (v*1.2.0*). The learning rate decayed by a factor of 10 every 20 epochs.

We determined the hyperparameters by exploring and testing different parameter settings with the validation dataset. Specifically, we explored four values (1, 5, 10, 15) for *β* and chose *β* = 10 to balance the reconstruction performance vs. the disentanglement (or independence) of latent variables (Figure 2) – the two terms in the loss function shown in Eq. (1). We also explored several options for the number of layers and the learning rate, and finalized those parameters based on the loss evaluated with the validation dataset (Supplementary Figure 2 and 3). Note that with the training strategies described above, only VAE model with 12 layers were able to reduce both reconstruction loss and *D*_*KL*_ when *β* = 10.

**Figure 2.**
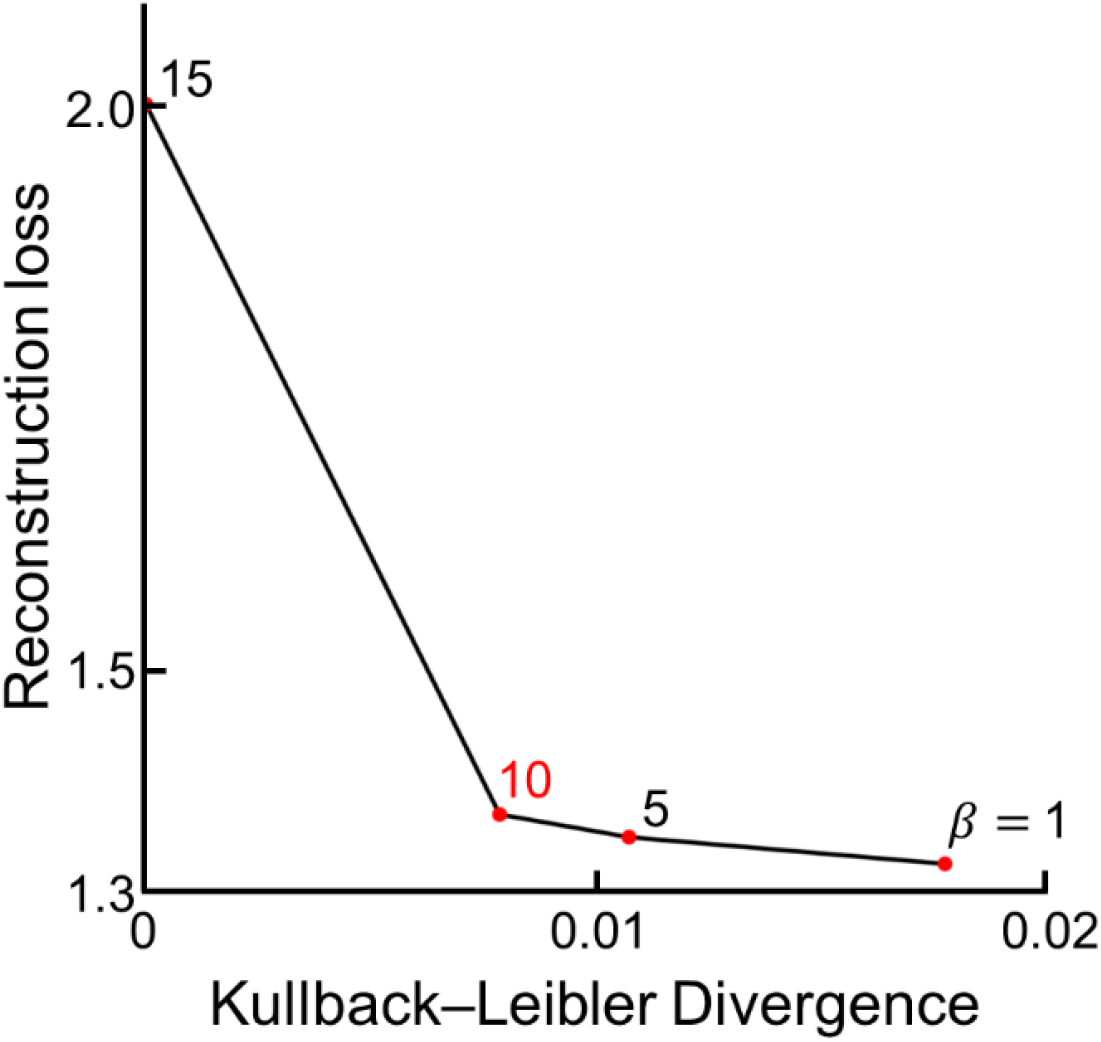
Validation losses for the VAE models with different values for *β* in Eq. (1). *β* = 10 shows a reasonable trade-off between the reconstruction loss and the Kullback–Leibler divergence on the validation dataset.

### Synthesizing rsfMRI functional connectivity

We used the trained VAE to synthesize rsfMRI data from random samples of latent variables. To synthesize a vector in the latent space, we drew a random sample of every latent variable independently from a standard normal distribution. The synthesized vector passed through the decoder in VAE, generating a cortical pattern. Repeating this process, we synthesized 12,000 cortical patterns as data used for seed-based correlation analysis. As examples, we explored three seed locations within V1, IPS, and PCC and calculated the functional connectivity to each seed based on the Pearson correlation coefficient. The MNI coordinates of the seed in V1, IPS, and PCC were (7, −83, 2), (26, −66, 48), and (0, 57, 27), respectively (Jarrett, 2009). In addition, we performed a similar analysis without limiting to the seed locations. Instead, we calculated the functional connectivity between each pair of parcels as defined in a 360-parcel atlas of the whole cortex (Glasser et al., 2016).

For comparison, we similarly calculated seed-based or parcel-to-parcel functional connectivity with experimental rsfMRI data concatenated across a varying number (1, 5, 10, 50, and 100) of subjects in HCP. We compared the functional connectivity pattern observed with synthesized and experimental data and repeated the comparison 20 times. At each time, we generated a different set of synthesized data while using experimental data from a different subset of subjects. The comparison was thus randomly repeated.

### Defining a principal basis set in the latent space

By our design, the VAE model encodes the spatial pattern of fMRI activity and does not represent the temporal dynamics explicitly. The distribution of every latent variable is constrained to be close to a standard normal distribution independent of one another, for the K-L divergence term in the loss function in Eq. (1). This implies that the latent variables in the VAE model are not unique. An arbitrary rotation of a tentative set of latent variables would arrive at a new set of latent variables that span the same latent space and satisfy the same learning objective.

To identify a unique set of latent variables, we exploited the trajectory of the latent representation. Specifically, for the fMRI data in the testing set (concatenated across 500 subjects), we encoded the fMRI pattern observed at every time into a point (or vector) embedded in the latent space. As time progressed, this point moved in the latent space along a trajectory that represented the temporal dynamics of fMRI activity.

In a first-order differential analysis, we evaluated the displacement (or difference) of the latent representation from every time point to its next. To this time-difference vector (or the latent gradient), we further applied singular vector decomposition and used the singular vectors to define a unique basis set of the latent space. As such, each singular vector was a re-defined latent variable, while the corresponding singular value indicated its importance in explaining the latent gradient of cortical activity. In other words, the trajectory was more likely to move along the direction represented by a singular vector with a larger singular value than that with a smaller singular value.

We further interpreted and visualized the top-10 latent variables defined as the singular vectors with the largest 10 singular values. For this purpose, we decoded each of these latent variables onto the cortical surface by using the decoder in the VAE model. Note that the latent variables were related to cortical patterns through nonlinear functions. We evenly sampled each latent variable of interest from −5 to 5, while keeping other latent variables to zero. We mapped the decoded cortical pattern and characterized its variation due to the variation of a single latent variable. We quantified the variation separately for each cortical location in terms of the standard deviation multiplied by a sign. The sign of standard deviation map was determined by measuring the sign of Pearson correlation coefficient between the decoded values of each cortical location and the samples of the given latent variable.

### Clustering in the latent space

We encoded the rsfMRI spatial pattern at every time point for 500 testing subjects, yielding 600,000 vectors in the latent space. We used k-means clustering with 1-cosine distance (based on “kmeans” in Matlab) to group those vectors to 21 clusters. The choice of k=21 was empirical but made intentionally to be consistent to a prior study with a similar motivation (Smith et al., 2012). This choice was within a reasonable range for the number of resting state networks as reported in literature (Smith et al., 2009; Yeo et al., 2011). Beyond this single choice, we explored other numbers of clusters to ensure that k=21 was a reasonable choice for the distribution of latent representation. Specifically, we varied k from 1 to 100. Given each choice, we ran the k-means clustering for the testing data (n=500 subjects), identified the clusters, calculated the centroid of each cluster, and summed the distance to the centroid within every cluster. We further plotted the sum of distance as a function of k and ensured that k=21 was around the “elbow” of the plot, as a useful, but not strict, rule of thumb.

Given k-means clustering with k=21, we re-ordered individual time points by their cluster membership and compared the distance between time points within and between clusters. We also evaluated the relationships among different clusters by calculating the distance between the centroids of individual clusters and we further grouped clusters into super-clusters organized in a multi-level hierarchy visualized as a dendrogram (by using “linkage” in Matlab).

To visualize and interpret each cluster, we further converted the cluster centroid to a corresponding cortical pattern by using the VAE’s decoder. The resulting cortical pattern was scaled such that its maximal absolute value equaled 1. This pattern was considered as a functional cortical network. To further evaluated how each cortical network changed its activity in time, we defined and evaluated the cluster-wise activity as the cosine affinity between the centroid of each cluster and the latent representation of fMRI activity at every time. As such, a cluster increased its activity when the latent representation moved toward the centroid of that cluster or decreased its activity when the representation moved away from that centroid but towards the centroid of another cluster. After this analysis was done separately for each session and subject, we averaged the cluster-wise activity across subjects. Then we compared the group-level activity between sessions and across clusters, and tested the statistical significance with a non-parametric permutation test (false discovery rate q<0.01), for which the time points were randomly shuffled for 10,000 trials to yield a null distribution.

### Individual variation

To evaluate the individual variation, we compared the latent representations of the fMRI data from different individuals. In an exploratory analysis, we randomly selected a small (n=20) subset of subjects. We chose 20 subjects to ease visualization and intuitive demonstration, before scaling up the analysis to 500 subjects. For each of the 20 subjects, we converted the fMRI activities, instance by instance, to the representations in the latent space. To visualize and compare subject-wise representations, we used the t-distributed Stochastic Neighbor Embedding (t-SNE) method to visualize the 256-dimensional latent representations (color-coded by subjects) in a two-dimensional space. We calculated the Silhouette index to measure how similar the latent representation was within the same subject vs. between different subjects.

### Individual identification

After the exploratory analysis above, we evaluated the individual variation across n=500 subjects. For the distribution of subject-wise latent representation, the first moment was the mean and the second moment was the covariance, which indicated the location and geometry of the subject-wise latent representation, respectively. We tested the use of the first moment (mean) or the second moment (covariance) as the subject-identifying feature.

In the testing data set, every individual had rsfMRI data acquired for two separate sessions. From the first session, we extracted the feature from every subject and stored it as the subject-identifying key in a database that included a population of 500 subjects. Given this database, we tested the accuracy of retrieving any subject’s identity by using the feature extracted from the second session as a query to match against all keys in the database. The goodness of match was evaluated as the cosine similarity or the Pearson correlation coefficient when the query and the key were based on the first moment (mean) or the second moment (covariance) of the subject-wise representation, respectively. The accuracy of individual identification was evaluated as the percentage by which the correct identity was retrieved as one of the best 1, 5, or 10 matches, yielding the namely top-1, 5, or 10 accuracy.

For comparison, we compared the performance of individual identification based on the above latent-space feature vs. the similar feature evaluated in the cortical space. The cortical-space features extracted with a similar method as previously reported in (Finn et al., 2015). Specifically, the FC between brain regions (or connectome) was calculated as features for individual identification. It is worth noting that the cortical connectome and covariance of latent representation, although they are nominally different terms, can both be viewed as the representational geometry of brain activity in the cortical space (for the connectome) or the latent space (for the covariance of latent representation). In addition, we may also cast both notions as the functional connectivity profile in the cortical space or the latent space. Given such conceptual connections, we evaluated the FC between every pair of 360 cortical parcels defined in an established atlas (Glasser et al., 2016) and used the FC-based connectome as the feature for individual identification (Finn et al., 2015). We compared the connectome-based identification accuracy with that based on the FC profile (or representational geometry) in the latent space for a varying population size (from n=5 to 500 subjects) or a varying length of data per subject (from 9 to 180 s). We repeated the above analysis 100 times, each time with a different subset of the testing data and averaged the identification accuracy across the repeated tests.

### Comparison with linear latent space

The VAE model described herein provided nonlinear mapping from the cortical space to the latent space (through the encoder) and in reverse (through the decoder). Such reversible mapping could be conventionally done through linear matrix operations, such as the principal component analysis (PCA) and independent component analysis (ICA). Hence, we compared the distribution and geometry of the rsfMRI representation in the nonlinear latent space obtained with VAE vs. the linear latent space obtained with PCA or ICA. For such comparison, we used PCA or ICA trained with the training data to represent rsfMRI data in the testing dataset, while keeping the linear latent space of the same (256) dimension as its nonlinear counterpart. We compared the performance of reconstructing fMRI patterns from their latent representations (see results in Fig. 4). In addition, we also compared PCA or ICA vs. VAE for characterizing individual variation or performing individual identification by using the representation in the PCA or ICA-derived linear latent space for the same analyses as used for the representation in the VAE-based nonlinear latent space (see results in Figs. 8 and 10).

## Results

### VAE compressed rsfMRI maps

Inspired by its success in artificial intelligence (Higgins et al., 2017; Kingma and Welling, 2013), we designed a VAE model in order to disentangle the generative factors underlying rsfMRI activity. The model was trained to represent and reconstruct rsfMRI data with a set of latent variables that were constrained to be as independent as possible. The hyper-parameter, *β*, which expressed the weighting of independence among latent variables in the overall learning objective, was initially explored for different values (1, 5, 10, 15) before being finalized to *β*=10 – a setting that led to a reasonable trade-off of the model performance vs. constraint as demonstrated with the validation dataset (Fig. 2).

The model used a pair of convolutional and deconvolutional neural networks in an encoder-decoder architecture (Figure 1.b). The encoder transformed any rsfMRI pattern, formatted as an image on a regular 2D grid (Figure 1.a), to the probability distributions of 256 latent variables. The decoder used samples of the latent variables to reconstruct or generate an fMRI map. Using data from HCP (WU-Minn HCP Quarter 2) (Van Essen et al., 2013), we first trained the model with rsfMRI maps from 100 subjects and then tested it with rsfMRI data from 500 other subjects.

After being trained, the model could compress any fMRI map to a low-dimensional latent space and restore the map from the latent representation separately for every time point (Figure 3). The compression resulted in spatial blurring comparable to the effect of spatial smoothing with 4-6 mm full width at half maximum (FWHM) (Figure 4). Given fMRI data spatially smoothed to a varying extent (FWHM from 1 to 10 mm), VAE showed either comparable or better performance of reconstruction than its linear counterparts (PCA and ICA), when VAE, PCA, and ICA all used the same dimension (256) for their latent spaces (Figure 4.a). The difference in reconstruction performance between VAE and PCA or ICA was marginal but statistically significant (repeated measures ANOVA followed by post-hoc paired t-test, false discovery rate *q*<0.05), for all smoothing levels except FWHM=1 mm (Figure 4.b). These results suggest that the latent representation obtained with VAE preserved the spatial and temporal characteristics of rsfMRI, despite a modest but acceptable loss in spatial resolution and specificity.

**Figure 3.**
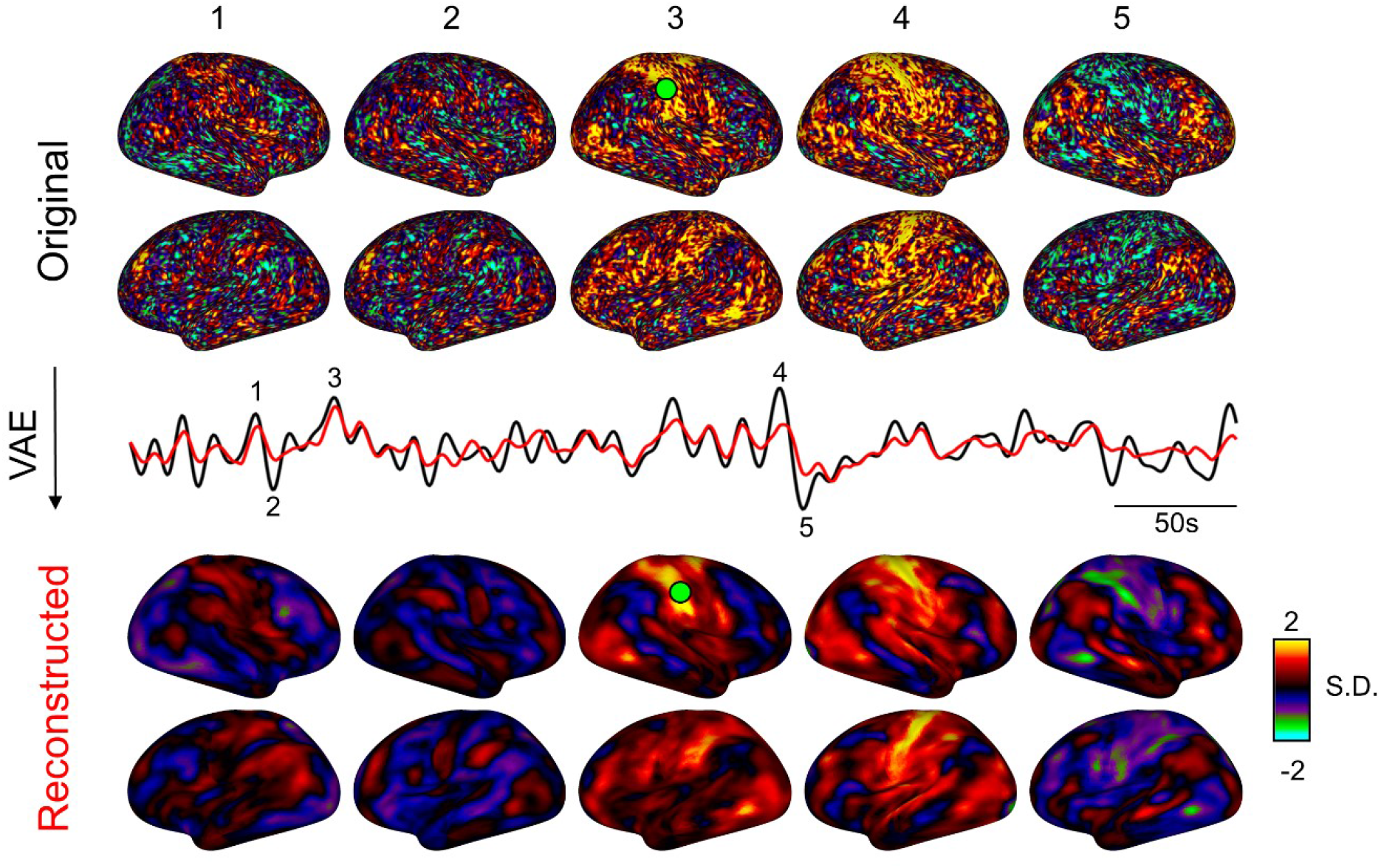
Image reconstruction using VAE. A series of cortical patterns are reconstructed through the VAE model. Among them, five original cortical patterns (upper panel) and their corresponding reconstruction through VAE (bottom panel) are visualized for comparison. For an example region (green circle), the time series of the original activity (black line) and the reconstructed activity (red line) are plotted for comparison.

**Figure 4.**
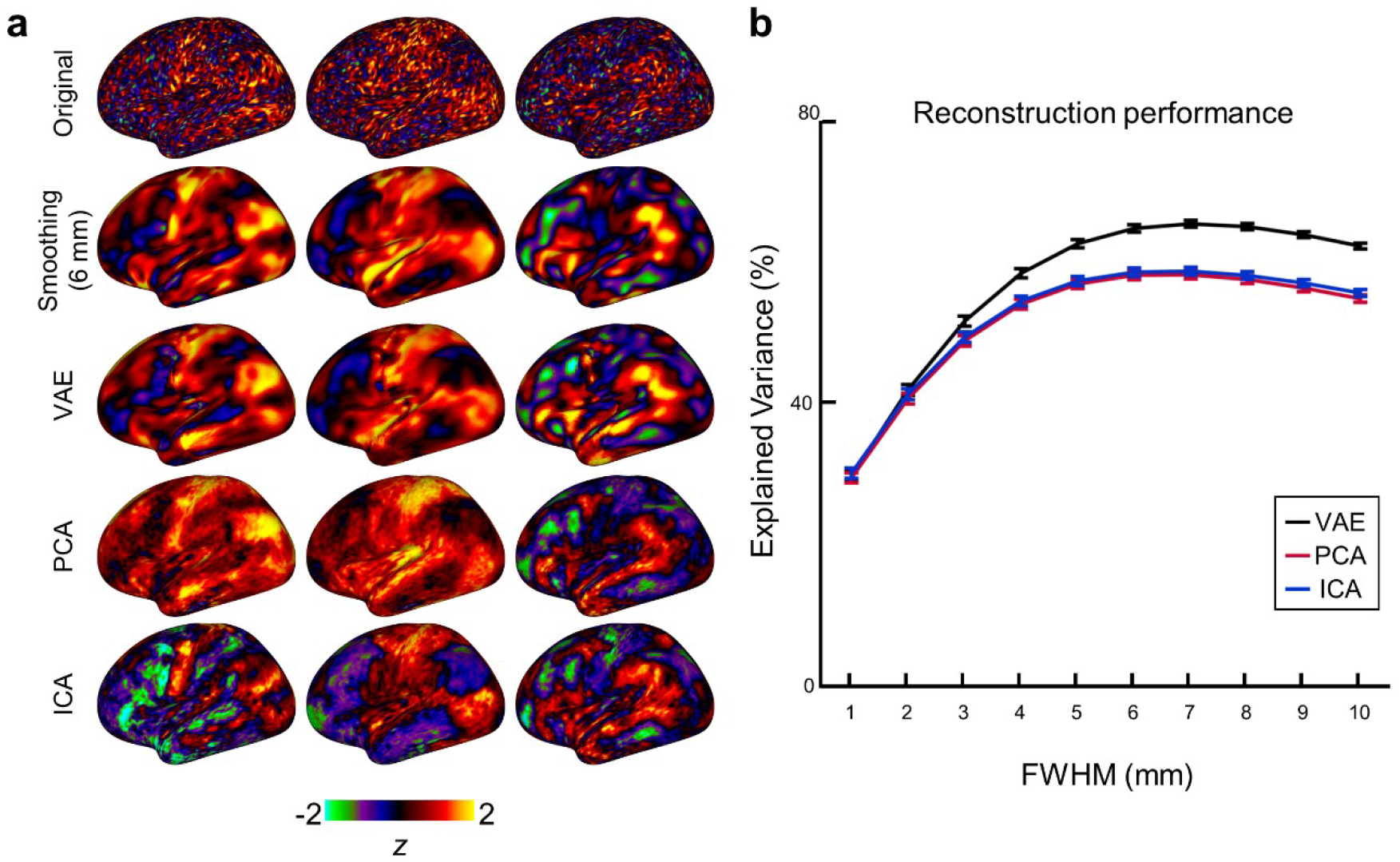
RsfMRI data compression and reconstruction with VAE vs. PCA and ICA. (**a**) For illustration, three example maps of fMRI activity, before (1^st^ row) and after (2^nd^ row) being smoothed (FWHM=6mm), are shown in comparison with the corresponding maps reconstructed with VAE (3^rd^ row), PCA (4^th^ row), and ICA (5^th^ row) trained to compress and reconstruct the training data from 100 subjects with 256 variables or components. (**b**) For quantitative comparison, the reconstruction performance, in terms of the percentage of variance in the fMRI images as explained by the model reconstruction, is shown for VAE, PCA, and ICA as a function of FWHM (from 1 to 10 mm) applied to the spatial smoothing of the fMRI images. The error bar stands for the standard error of mean.

### VAE synthesized correlated fMRI activity

We asked whether the decoder in the VAE, as a generative model of fMRI activity, had learned the putative mechanisms by which rsfMRI activity patterns arise from brain networks. To address this question, we randomly sampled every latent variable from a standard normal distribution and used the decoder to synthesize 12,000 rsfMRI maps (equivalent to 10 subjects at 1,200 time points per subject).

We calculated the seed-based correlations by using the VAE-synthesized data and compared the resulting maps of correlations with those obtained with rsfMRI data concatenated across a different number of subjects. Figure 5.a shows three examples with the seed region in the primary visual cortex (V1), intraparietal sulcus (IPS), or posterior cingulate cortex (PCC). For each of the three seed locations, the synthesized fMRI data showed a similar correlational map as that based on length-matched rsfMRI data obtained from 10 subjects (Figure 5.a), and the correlational map was consistent with the literature (Yeo et al., 2011). The measured FC patterns were more similar to the synthesized FC patterns, when the measured FC was based on data from increasingly more subjects, regardless of whether the FC was evaluated and compared with respect to a specific seed location (Figure 5.b) or across all cortical parcels (Figure 5.c). These results suggest that the VAE provided a computational account for the generative process of resting state activity and could synthesize realistic rsfMRI activity patterns and preserve inter-regional correlations as are experimentally observable at a group or population level. However, it is worth mentioning that the temporal ordering of the synthesized data is not meaningful, since the VAE model does not explicitly model the temporal dynamics.

**Figure 5.**
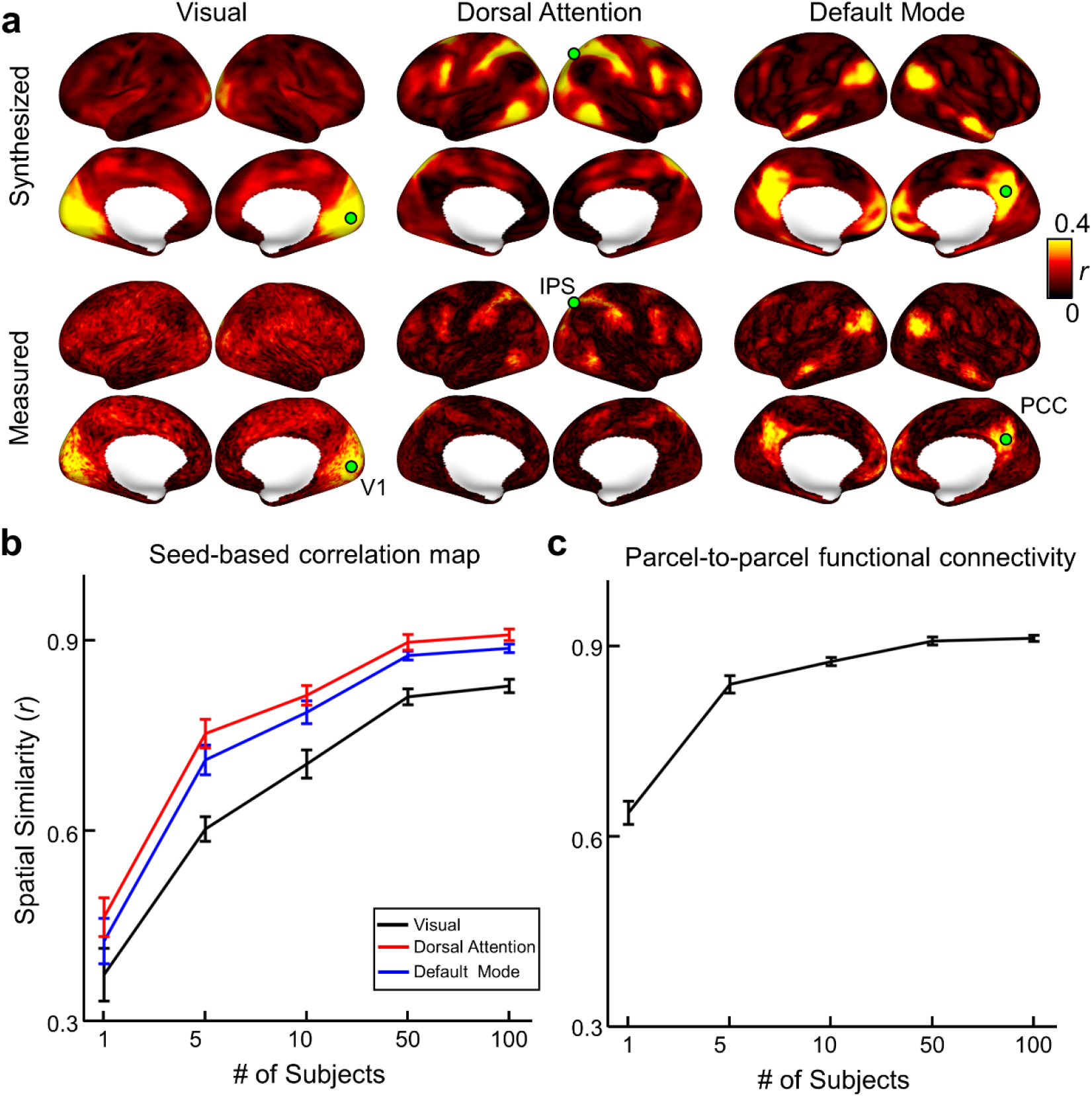
VAE synthesizes correlated fMRI activity. (**a**) Seed-based correlations of VAE-synthesized fMRI data (top row) vs. experimental fMRI data (bottom row) with the seed location (green circle) at V1 (left), IPS (middle), or PCC (right). (**b**) Spatial correlations between the seed-based functional connectivity based on VAE-synthesized data and those based on measured fMRI data concatenated across 1, 5, 10, 50, or 100 subjects. The colors indicate different seed locations (V1: black; IPS: red; PCC: blue). Similarly, (**c**) shows the spatial correlation between the synthesized vs. measured functional connectivity among 360 cortical parcels. The error bar indicates the standard error of the mean averaged across 20 repeated trials.

### Latent variables reflected network dynamics

We also examined the time-evolving trajectory of the latent representation and re-defined the latent variables such that they reflected the dynamic changes of fMRI activity. We first evaluated the displacement of the latent representation from every time point to its next. Then we applied singular value decomposition and used the resulting singular vectors to redefine the latent variables as a new basis set that spanned the latent space. These redefined latent variables, ranked in a descending order by their singular values, represented the principal directions in which the instantaneous latent representation tended to move along its time-evolving trajectory.

We chose the top-10 latent variables for further visualization and interpretation. For each latent variable, we uniformly sampled its value in a range from −5 to 5 and visualized each sample by decoding it to a cortical pattern. We found that as the latent variable increased its value linearly, the decoded cortical pattern changed in a non-linear way that differed across cortical locations (see illustrative examples in Figure 6.b). To visualize how each latent variable controlled the activity at each cortical location, we calculated the standard deviation of the voxel-wise activity change given an increasing value for the given latent variable and multiplied by a sign (+1 or −1) depending on whether the activity tended to increase or decrease as the latent variable increased. For example, the 1^st^ latent variable was visualized as a cortical pattern that resembled the default mode network (Figure 6.a). Using the same visualization method, we found that the 2^nd^ through 10^th^ latent variables all corresponded to distinct but partially overlapping cortical patterns (Figure 6.d). However, the top-10 latent variables were found to be inadequate to explain the dynamics of latent representations. The percentage of the variance explained by each latent variable was around 1% or less, and the total variance collectively explained by top 10, 20, 50, and 100 latent variables were 9.6, 17.5, 37.6, and 62.3% (Figure 6.c). These results suggest that the dynamics of rsfMRI is complex and high-dimensional in nature. Nevertheless, the latent variables derived from the above analysis represent distinctive factors that drive the dynamic change in resting state activity.

**Figure 6.**
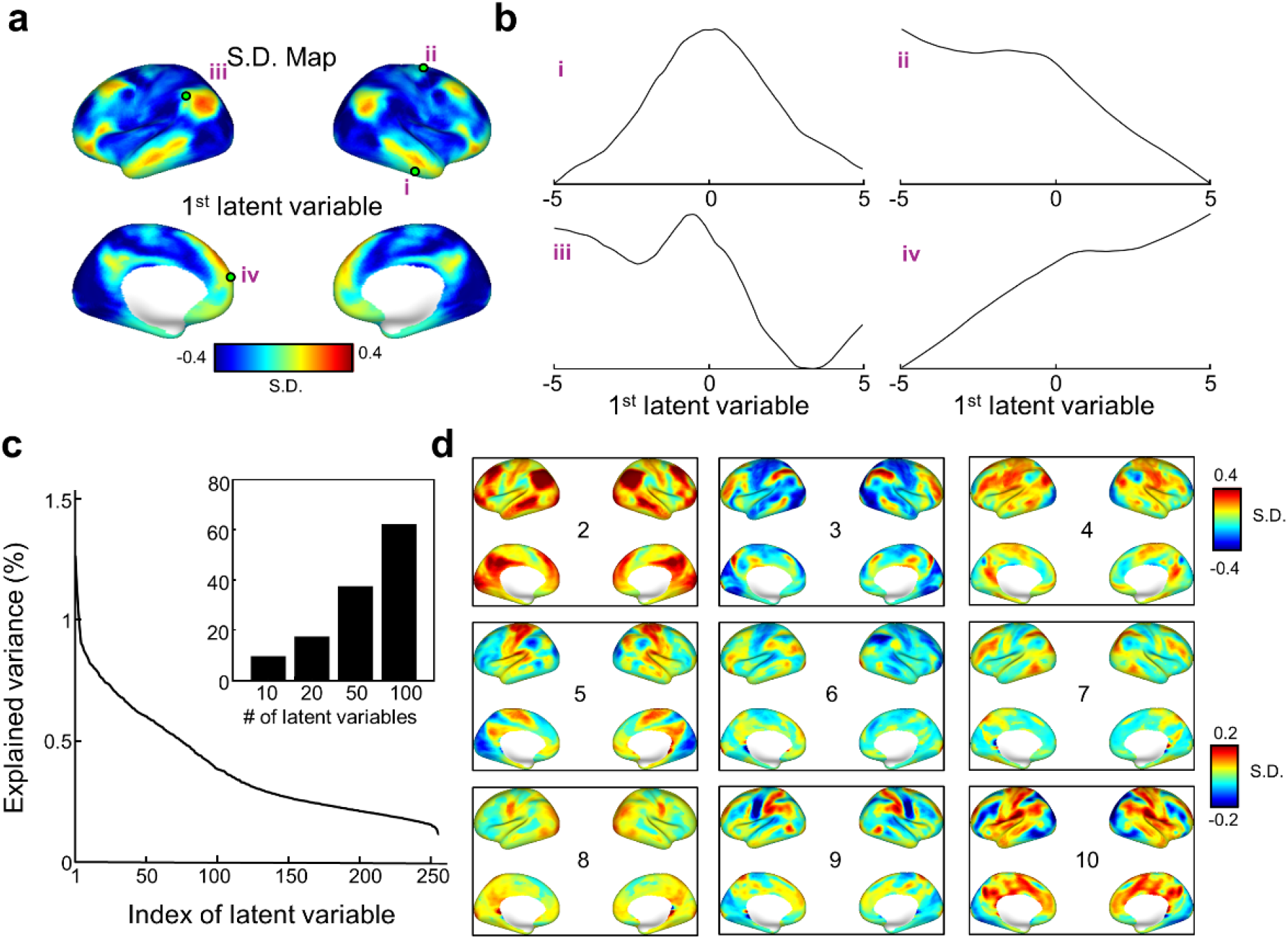
Latent variables drive the dynamics of latent representation. The latent variables are defined as the principal directions for the dynamic change of latent representation across adjacent time points. **(a)** Visualization of the 1^st^ latent variable as a cortical pattern of the signed standard deviation. (**b**) For each of the four cortical locations, denoted as i through iv and shown as green circles in (**a**), the activity change is shown as a function of the 1^st^ latent variable, indicating a varying nonlinear relationship. (**c**) The percentage of the variance that each latent variable explains the first order dynamics of latent representation. The inset shows the percentage of the total variance explained by top 10, 20, 50, or 100 latent variables. (**d**) The visualization of the 2^nd^ through 10^th^ latent variables as cortical patterns.

### Clusters in latent space

We further characterized the distribution of latent representation and attempted to identify clusters in the latent space. We used the VAE to encode the rsfMRI pattern observed at every time point from 500 subjects, clustered the time points by applying k-means clustering (k=21) to the latent representations, and decoded the cluster centroids to corresponding cortical maps. The number of clusters (k=21) was close to the “elbow” indicative of a reasonable balance between reducing variation within clusters and avoiding too many clusters (Figure 7.a). The 1-cosine dissimilarity between latent representations reordered by their cluster membership shows not only close affinity within every cluster but also a varying level of affinity between different clusters (Figure 7.b). This motivated us to hierarchically merge clusters into super-clusters (or “clusters of clusters”) based on the cosine affinity between cluster centroids (Figure 7.c).

**Figure 7.**
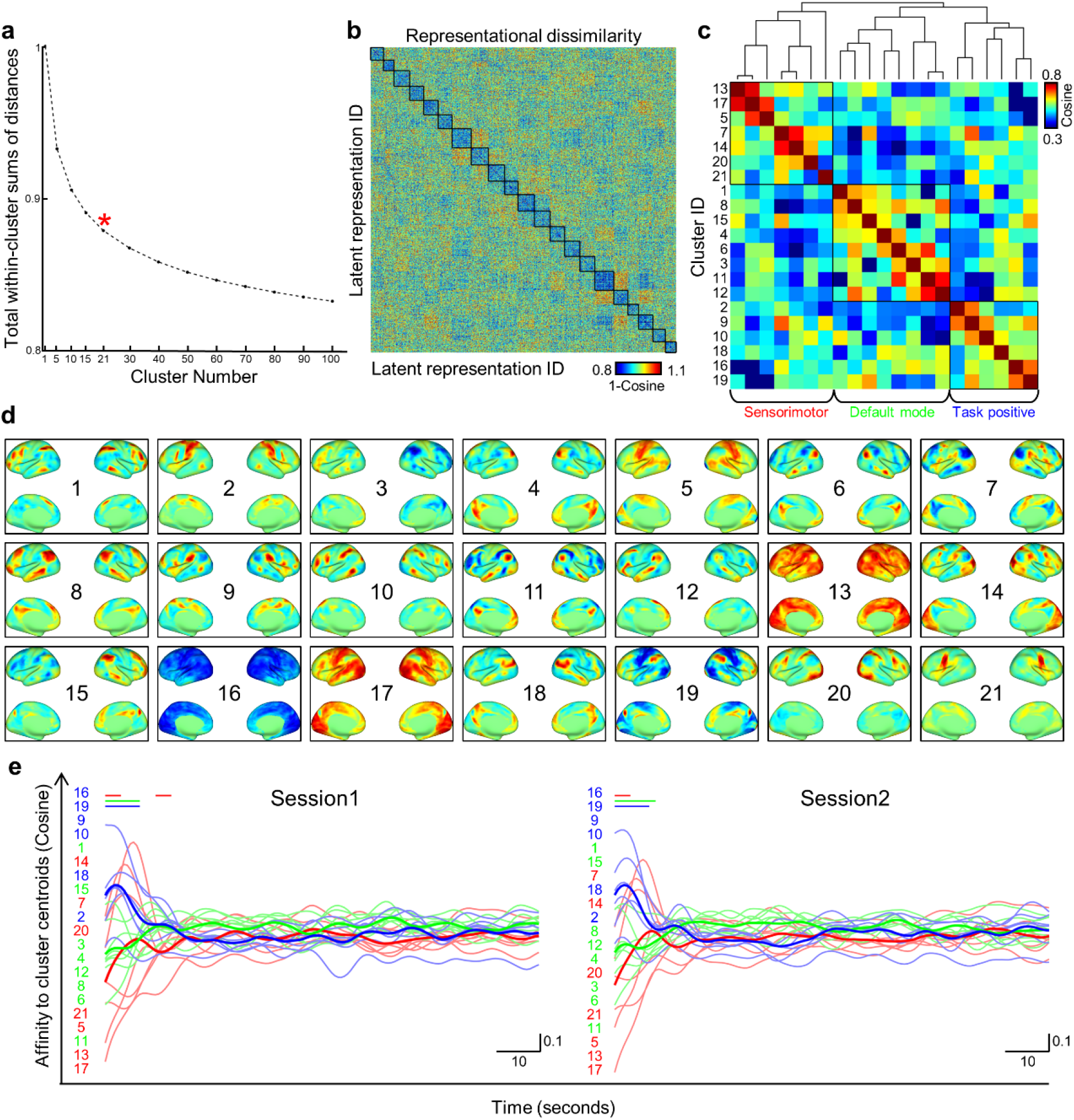
Clusters of latent representations. (**a**) The sum of within-cluster distance is shown as a function of the number of clusters (or K). The choice of k=21 is about where an elbow is observable in the plot. (**b**) Representational dissimilarity matrix of the 1-cosine distance between instantaneous latent representations (from 100 subjects with 1,200 time points per subject) reordered by cluster membership. (**c**) Hierarchical clustering of 21 cluster centroids. The clusters are grouped into three super-clusters, provisionally labeled as sensorimotor (red), default mode (green), and task positive (blue) networks, based on the cortical visualization of individual clusters in (**d**). (**d**) Cortical patterns decoded from every cluster centroid. The number shows the cluster index. The scale of each pattern is normalized by its maximal absolute value. (**e**) The group-averaged cluster-wise activity, described as the cosine affinity of instantaneous representation to the centroid of each cluster. Each thin line corresponds to one cluster; the color indicating the super-cluster (sensorimotor: red; default mode: green; task positive: blue) that each cluster belongs to. The thick lines show the average within super-clusters. The left and right panels show the activity patterns averaged across all subjects for session 1 and session 2, respectively. Colored lines on the top of panel **(e)** highlight the periods in which the sensorimotor (red), default-mode (green) or task positive (blue) super-cluster was statistically significant in the group level (permutation test, false discovery rate q<0.01).

For each of the 21 clusters, we decoded and visualized the cluster centroid as a cortical pattern as shown in Figure 7.d. Among the 21 clusters, 5 clusters (Cluster 4, 6, 11, 12, 18) showed activity increase (positive) at one or multiple regions in the default mode network (Buckner et al., 2008; Greicius et al., 2003; Raichle et al., 2001), alongside activity decrease (negative) at other regions. Similarly, we found 5 clusters with activity increase in the so-called frontoparietal control network (Cluster 8) (Dixon et al., 2018), cingulo-opercular network (Cluster 7 and 9) (Dosenbach et al., 2007), cognitive control network (Cluster 1) (Cole and Schneider, 2007), and dorsal attention networks (Cluster 10) (Fox et al., 2006) – collectively referred to as “the task positive network” (Fox et al., 2005). In addition, cluster 13 and 16 showed activity decrease in the whole brain, thereby a signature of global signal fluctuation (Murphy et al., 2009; Schölvinck et al., 2010; Wen and Liu, 2016). Cluster 5 and 17 showed widespread synchrony across sensory systems. Cluster 2 and 21 showed the networks for sensorimotor control of the limbs and of the mouth, pharynx, and visceral organs, respectively. Whereas most clusters were bilaterally symmetric, Cluster 15 and 3 were unilateral to the right and left prefrontal cortex, respectively. A common observation for many clusters was that a cluster could highlight the positive interactions among a set of well-defined cortical regions alongside their negative interactions with a different set of regions. Given the above interpretation of individual clusters, we further interpreted the three super-clusters as sensorimotor, default mode, and task positive networks (Figure 7.C).

In addition, we evaluated the temporal dynamics of latent representation in terms of the dynamics of individual clusters or their corresponding cortical networks. Intuitively, we considered the time-evolving trajectory of the latent representation as the movement towards or away from each cluster. In this regard, we expressed the cluster-wise activity as the time series of cosine affinity between the instantaneous latent representation and the cluster centroid. During a rsfMRI session, different clusters expressed similar activity levels (Figure 7.e), except in the initial period of the session. In that period of 20 seconds, clusters presumably related to task positive networks showed a transition from a high activity level to a lower steady state; the clusters related to sensorimotor networks showed a transition from a low activity level to a higher steady state; in contrary, clusters related to the default mode network remained roughly unchanged. These (somewhat incidental) observations were consistent and reproducible across individuals and sessions. On one hand, this result suggests that the first 20 seconds in a rsfMRI session are not necessarily the steady state under a resting condition. On the other hand, this exploratory analysis shows the feasibility of using the VAE-extracted latent representations to identify brain networks and reveal their individual dynamics.

### Individual variation of latent representation

Whereas the aforementioned analyses focused on the group-level characteristics of the latent representations, we further asked how the distribution and geometry of latent representation varied across individuals. Only for the sake of demonstration, we randomly selected 20 subjects in the testing dataset and visualized their individual representations in the latent space after reducing its dimension from 256 to 2 by using t-SNE (Figure 8.a). Strikingly, the latent representations were grouped by and separable across individuals. The clustering by individuals was noticeable in the nonlinear latent space obtained with VAE (Figure 8.a), but not in the linear latent space obtained with PCA (Figure 8.b). Such distinctions were quantitatively confirmed (Figure 8.c) by using the Silhouette value to measure the degree of clustering by individuals. The Silhouette value for VAE (mean ± std: *s* = 0.044±0.002, 50 bootstrapping trials) was significantly higher (p<0.001, two-sample t-test) than that for PCA (*s* = −0.020±0.015). Using the center of latent representation as the subject-identifying feature, we found that subject identity could be retrieved with a reasonably high accuracy when the latent representation was extracted by VAE, whereas the linear representation by PCA failed the same task nearly entirely (Figure 8.d). These results suggest the feasibility of using VAE to characterize and reveal individual variations of resting state activity in a non-linear latent space.

**Figure 8.**
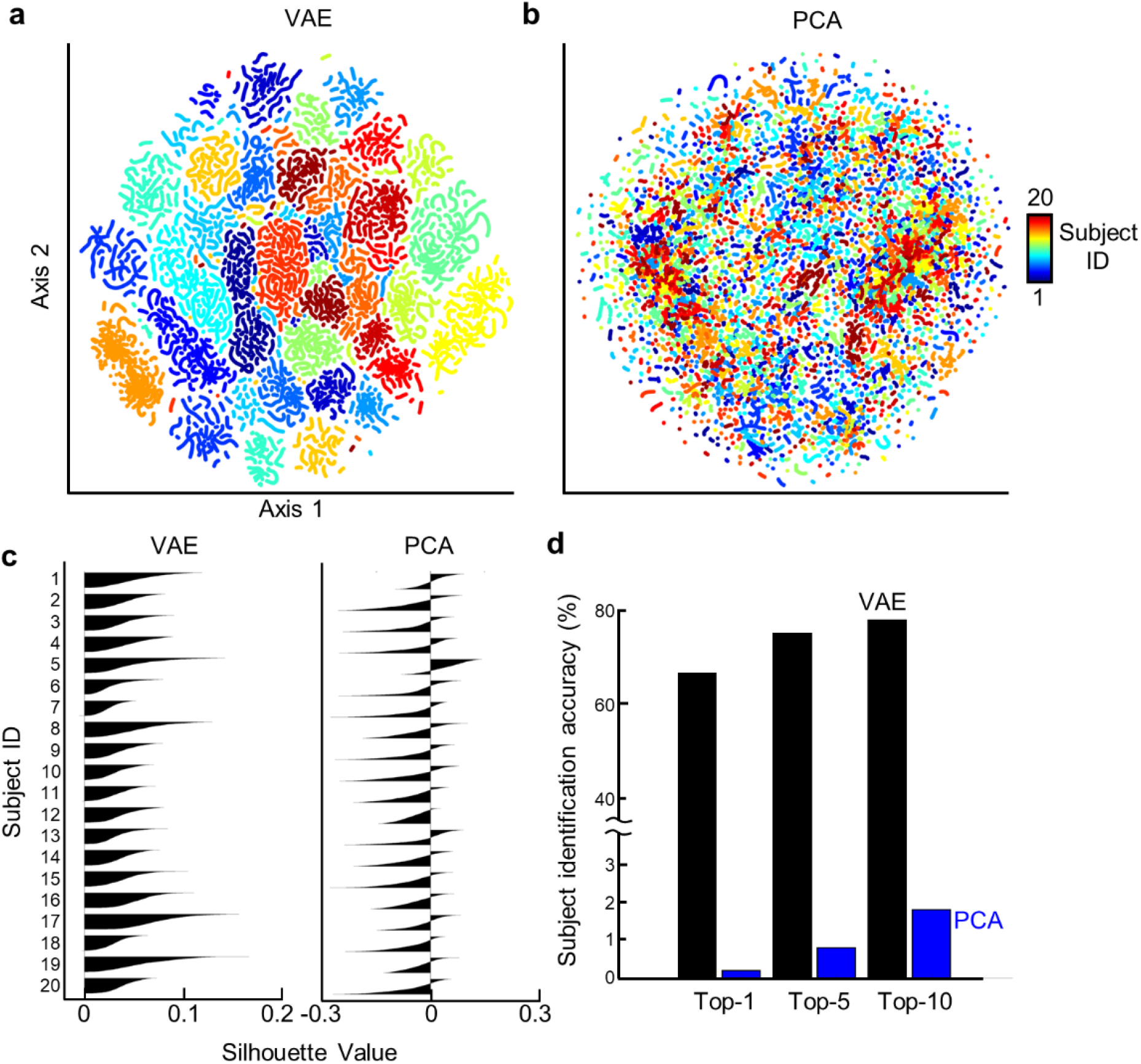
Individual variation of latent representation obtained with VAE vs. PCA. **(a-b)** Subject-wise latent representations visualized in a 2-D space obtained with t-SNE, when **(a)** VAE or **(b)** PCA is used to extract representations of rsfMRI activity from 20 subjects. **(c)** The Silhouette value shows how similar a representation is similar to each other within the same subject as opposed to between different subjects for VAE (left) or PCA (right). **(d)** The top-1, 5, and 10 accuracy of using the time-averaged representation as the feature to identify individuals in a large group of (n=500) subjects, for the representations obtained with VAE (black) or PCA (blue).

### Individual identification

From the t-SNE based visualization (Fig. 8.a), it was noticeable that subject-wise representations exhibited different geometries. Some were more elongated or scattered than others. This observation motivated us to ask whether the representational geometry (Kriegeskorte and Kievit, 2013) could be an individual-specific feature (or “fingerprint”) to allow for more accurate individual identification. Specifically, we calculated the covariance between every pair of latent variables and assembled the pair-wise covariance into a vector as the feature of the representational geometry and evaluated the similarity in this feature between two sessions within or between subjects. The representational geometry evaluated in this way could be interpreted as the functional connectivity (FC) between latent variables. This interpretation related this approach to a conceptually similar approach: the “connectome-based fingerprinting” (Finn et al., 2015; Venkatesh et al., 2020), in which the functional connectivity was evaluated between cortical parcels. So, we evaluated the use of either the latent-space or cortical-space FC for individual identification in comparison.

As shown in Figure 9.a, FC between any pair of cortical areas was mostly positive (mean ± std of z-transformed correlation: *z*=0.26±0.3) and highly reproducible not only within the same subject (*r*=0.66) but also between different subjects (*r*=0.45). On the other hand, FC between latent variables had both positive and negative values (mean ± std of covariance: *σ*^2^=0.00±0.13) and its reproducibility was high only within the same subject (*r*=0.33) but not between different subjects (*r*=0.07). The FC profile was more distinctive across subjects when it was evaluated between latent variables rather than cortical areas (Figure 9.b). In the latent space, the FC profile was significantly more consistent within a subject than between subjects (two-sample t-test, *t*(249,998)=254.05, two-sided *p*<0.001). The distribution of within-subject correlations was in nearly complete separation from that of between-subject correlations (Figure 9.b, bottom).

**Figure 9.**
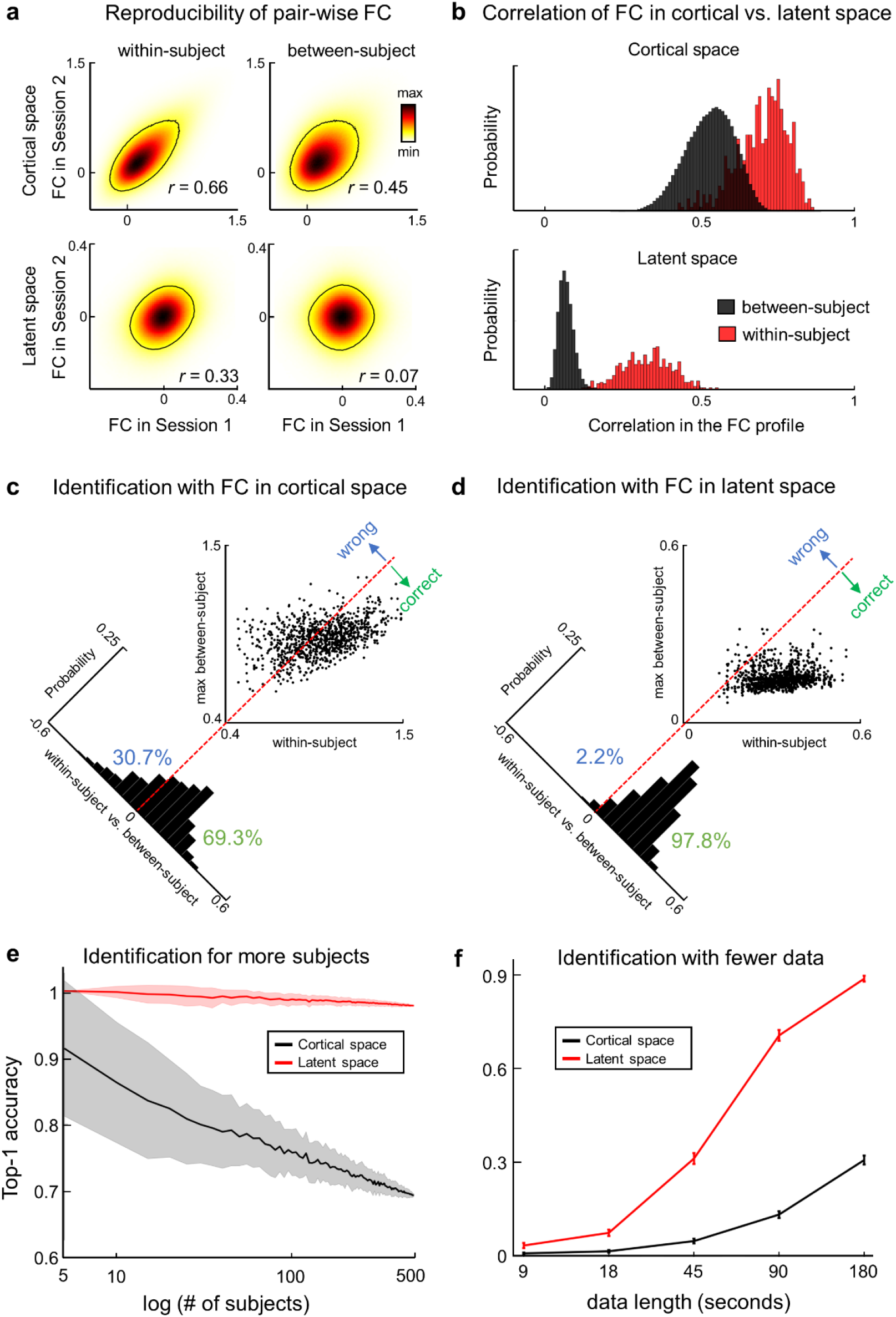
Individual identification based on correlations between latent variables or cortical parcels. **(a)** Density distributions of z-transformed correlations between every pair of cortical parcels (top) or covariance between every pair of latent variables (bottom). For each pair, the correlation and covariance in one session is plotted against the corresponding correlation in the other session for the same subject (within-subject, left) or different subjects (between-subject, right) given the testing dataset with n=500 subjects. Contour line stands for 20% of the maximal density. **(b)** Within-subject (red) and between-subject (black) correlations in the FC among cortical parcels (top) or latent variables (bottom) are shown as histograms with the width of each bin at 0.01. **(c)** In the scatter plot, each dot indicates one subject, plotting the maximal correlation in the cortical FC profile between that subject and a different subject against the corresponding correlation within that subject. The red-dashed line indicates y=x, serving as a decision boundary, across which identification is correct (x>y) or wrong (y>x). The histogram shows the distribution of y-x (0.05 bin width) with the decision boundary corresponding to 0. Similarly, (**d**) presents the results obtained with latent-space FC in the same format as (**c**). (**e**) Top-1 identification accuracy evaluated with an increasing number of subjects (n=5 to 500) given the latent-space (red) or cortical-space (black) FC profile. The solid line and the shade indicate the mean and the standard deviation of the results with different testing data. (**f**) Top-1 identification accuracy given rsfMRI data of different lengths (from 9s to 180s). The line and the error bar indicate the mean and the standard deviation with different testing data.

Then we compared the performance of individual identification on the basis of the FC profile in the latent vs. cortical space. To identify 1 out of 500 subjects, we compared a target subject’s FC profile in the 1^st^ session (as a query) against every subject’s FC profile in the 2^nd^ session (as a key) and chose the best match between the query and the key in terms of the Pearson correlation coefficient. As such, the choice was correct if the correlation with the target subject was higher than the largest correlation with any non-target subject. We found that the FC profile in the cortical space could support 69.3% top-1 accuracy while identification was often made with marginal confidence relative to the decision boundary (Figure 9.c). Using the FC in the latent space allowed us to reach 97.8% top-1 accuracy. The evidence for correct identification was apparent with a large margin from the decision boundary (Figure 9.d). The use of FC in the latent space supported reliable and robust performance in top-1 identification given an increasingly larger population (Figure 9.e) or when the data were limited to a short duration (Figure 9.f), being notably superior to the use of FC in the cortical space.

We further tested to what extent the performance of individual identification relied on the use of ICA-FIX to preprocess and denoise the rsfMRI data. For this purpose, we applied ICA-FIX to one or both of the two sessions in every subject and then tested the individual identification with n=500 subjects. As shown in Table 1, when the FC profile in the latent space was derived from the (ICA-FIX denoised) clean data for both the keys and queries, the identification has the highest accuracy (97.5%). When the key and the query were both based on noisy data (without denoising), the accuracy dropped to 91.3%. When the key and the query were unpaired as denoising applied to one but not the other, the accuracy further dropped to about 88%. Nevertheless, this performance obtained with the latent-space FC was still notably higher than the performance based on the cortical-space FC. For the latter, the use of unpaired preprocessing for the query and the key significantly dropped the identification performance from 69.3% to 47.5%. Counter-intuitively, when the denoising was applied to neither the query nor the key, the identification accuracy with the cortical-space FC increased to 76.9%, but still significantly lower than the accuracy of 91.3% obtained with the latent-space FC.

**Table 1.** Subject identification accuracy across different conditions

|  |  |  | Session 1 |  |
| --- | --- | --- | --- | --- |
|  |  |  | Clean | Noisy |
| Session 2 | Latent Space | Clean | 97.8% | 87.9% |
|  |  | Noisy | 88.4% | 91.3% |
|  | Cortical Space | Clean | 69.3% | 47.2% |
|  |  | Noisy | 47.5% | 76.9% |

Lastly, we explored whether the representational geometry (based on the profile of the covariance between latent variables) would yield a similar level of distinction across individuals for a linear latent space obtained with PCA or ICA. As shown in Figure. 10, PCA or ICA was not as effective as VAE. The top-1 accuracy of individual identification was 61.1% for PCA, 63.6% for ICA, in contrast to 97.8% for VAE. The within-subject vs. between-subject similarity in the geometry of linear representation obtained with PCA or ICA exhibited largely overlapping distributions, whereas the corresponding distributions were separated nearly completely for the nonlinear representations obtained with VAE.

**Figure 10.**
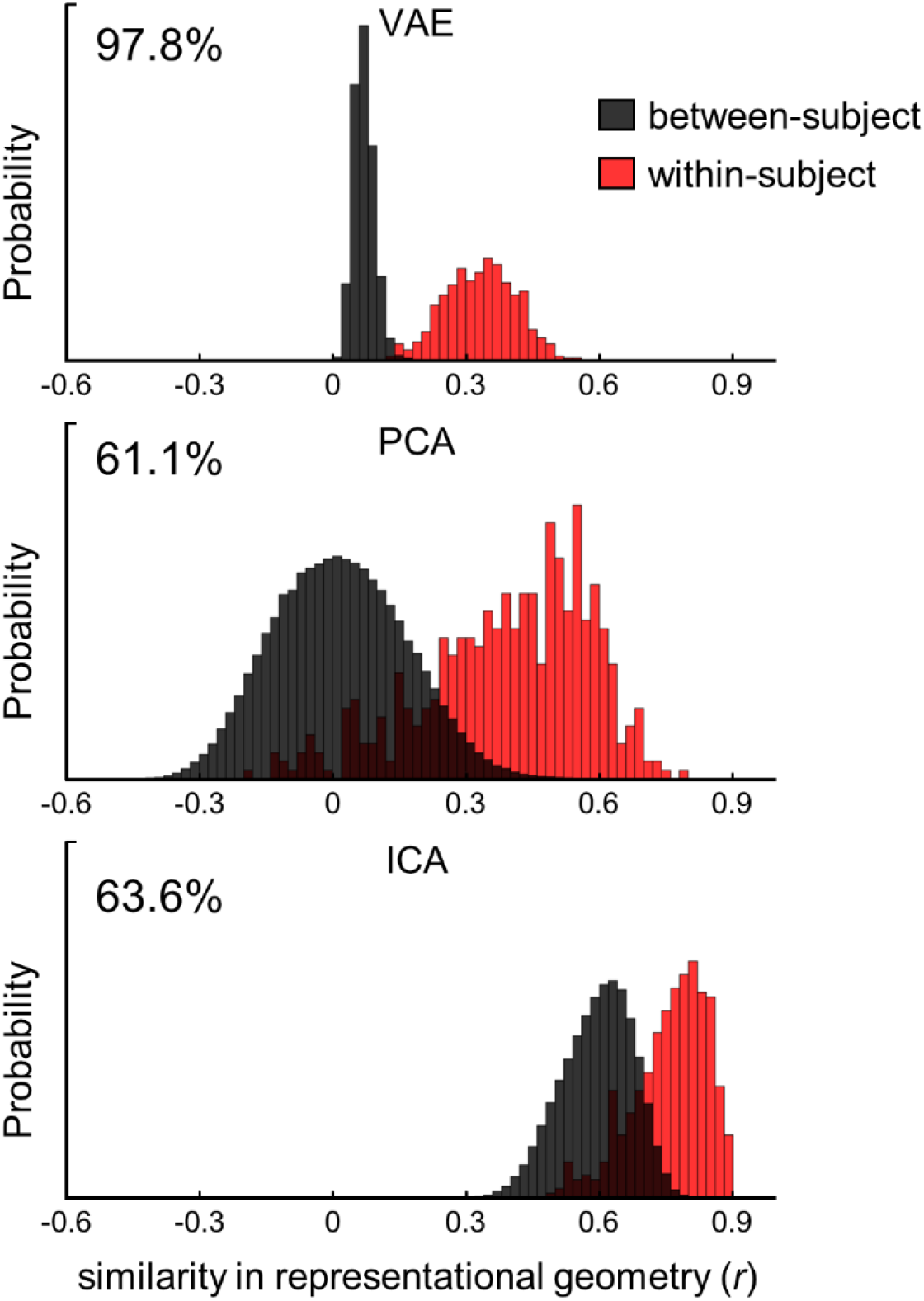
Individual identification with nonlinear vs. linear representations. Each plot shows the histogram of the similarity in the representational geometry between sessions within the same subject (red) vs. across different subjects (black), for representations in the nonlinear latent space obtained by VAE (top) or in the linear latent space obtained by PCA (middle) or ICA (bottom). The similarity reported is based on the inter-session correlation coefficient (or *r*). The histogram is discretized by bins with a width of 0.02.

## Discussion

Here, we present a method for unsupervised representation learning of cortical rsfMRI activity. Our results suggest that this method is able to disentangle generative factors underlying spontaneous brain activity, discover overlapping brain networks, capture individual characteristics or variation, and support accurate individual identification. We expect this method to be a valuable addition to the existing tools for investigating the origins of resting state activity, mapping functional brain networks, and potentially supporting individualized prediction of disease phenotypes and progression. Next, we discuss our findings from the joint perspective of methodology, neuroscience, and applications.

VAE is trainable with unsupervised learning (without any label) (Higgins et al., 2017; Kingma and Welling, 2013), which is appealing for learning representations of rsfMRI data. Since rsfMRI measures spontaneous brain activity unconstrained by any task, labels as required for supervised learning are either unavailable or far fewer than the data itself. Unsupervised learning with VAE can leverage the ever-increasing amount of rsfMRI data (Van Essen et al., 2013). The latent representations extracted from VAE can serve as the input to other algorithms to further support more specific goals such as classification of brain disorders and prediction of their phenotypes (Garrity et al., 2007; Moradi et al., 2015; Shen et al., 2010; Zhang et al., 2011).

The method herein can be extended in multiple ways. Although it is trained with rsfMRI data, we hypothesize that the VAE model can encode and decode both rsfMRI and task-fMRI data but with different latent distributions. If this is true, one may use this model to classify different perceptual, behavioral, or cognitive states and to reveal the distinctive network interactions underlying various states (Gonzalez-Castillo et al., 2015). The fact that the VAE can synthesize new data (Figure 5) is also appealing. It can be used as a post-processing strategy for data augmentation and interpolation, when data is short or corrupted, of interest for evaluation of dynamic functional connectivity (Allen et al., 2014; Chang and Glover, 2010) and correction for head motion (Power et al., 2014). It also supports the notion that the learned latent space captures the origins of rsfMRI and the VAE decoder captures the computational account for how rsfMRI arises from its origins.

It is worth mentioning two limitations of the VAE model in its current form. First, the model focuses on cortical patterns but excludes sub-cortical and white-matter voxels. This design is not only for the ease of model implementation but also for the predominant role of the neocortex in brain functions (Rakic, 2009). However, this precludes the model from accounting for subcortical networks or their interactions with the cortex. Addressing this limitation awaits future studies to redesign the model as a 3-D neural network that takes volumetric fMRI data as the input. Second, the VAE model only represents spatial patterns but ignores temporal dynamics inherent to rsfMRI data. Modeling the temporal dynamics is desirable but non-trivial, since it is highly irregular, complex and variable. To fill this gap, we direct future studies to designing a recurrent neural network (Chen and Hu, 2018; Cui et al., 2019; Shi et al., 2018; Sutskever et al., 2014; Zhao et al., 2019), as an add-on to VAE, to further learn sequence representation, for example, with a self-supervised predictive learning strategy (Kashyap and Keilholz, 2020; Khosla et al., 2019a).

Although VAE does not explicitly model the temporal dynamics, the representation obtained with VAE preserves the temporal dynamics (Figure 3). The trajectory of the latent representation describes the temporal behavior of brain networks, as opposed to voxels or regions. This trajectory is amenable to the use of many methods previously described for voxel-wise or region-wise analysis. To note a few examples explored in this study, the first-order temporal difference in the latent representation captures the gradient of latent trajectory that drives the brain to change its activity pattern from one time point to the next. As the latent gradient is also represented as a vector in the latent space, the length of this vector measures the displacement in the latent space and presumably the magnitude of network activity, and the direction of this vector encodes a pattern of network interaction that drives the instantaneous change of brain activity. The principal components of the displacement in representation uncover the important hidden factors that drive the temporal dynamics of brain networks (Figure 6). Similar analysis or notion has also been explored in two independent studies discussed in two very recent papers published or in preprint during the peer review of our paper (Brown et al., 2020; Liu et al., 2020b). These initial analyses are expected to merit and direct future studies upon predictive modeling of the trajectory of the VAE-derived latent representation, for example, by using Multivariate Auto-Regressive models (Liégeois et al., 2019; Rogers et al., 2010), Hidden Markov Models (Eavani et al., 2013; Suk et al., 2016).

VAE provides a new tool for mapping overlapping functional networks in the brain. A brain region may be involved in multiple networks each supporting a distinctive function (Liu and Duyn, 2013; Smith et al., 2012). However, existing network analyses still tend to group brain regions into non-overlapping networks (Yeo et al., 2011). VAE allows us to discover overlapping networks as clusters in the latent space spanned by independent latent variables. As such, VAE is conceptually similar to temporal ICA (Smith et al., 2012) but allows for nonlinear relationships between latent variables and the input data they represent (Khemakhem et al., 2019). Arguably, finding clusters in the low-dimensional latent space is more desirable than doing so in the higher-dimensional voxel space (Liu et al., 2013). Not only is it more computationally efficient, but representations are also more disentangled in the latent space than in the voxel space to readily reveal the underlying organization. However, it is not readily straightforward to attribute a cluster in the latent space to a distinct brain state (Hutchison et al., 2013) or an individual (Xie et al., 2018). Both are plausible. Our results show that individual variation manifests itself as the latent representation is in part clustered by subject (Figure 8), suggesting individual variation is a contributing factor to the clustering of latent representation. Our results also suggest that cluster-wise activity shows a consistent pattern across all subjects, in particular for the first 20 seconds of each session (Figure 7). Moreover, the clusters seem to group themselves hierarchically into presumably functional domains: sensorimotor, default-mode and task-positive networks (Figure 7). Together These results lead us to speculate that variation in brain states and individuals both contribute to the clustering of brain activity in the latent space. It is challenging to fully separate them and awaits future studies.

Central to this study is the efficacy of using VAE to disentangle what causes resting state activity. In the VAE model, the sources are the latent variables; the decoder describes how the sources generate the observed activity; the encoder models the inverse inference of the sources from the activity. Since the latent variables are data-driven, it is currently unclear how to interpret them as specific physiological processes, many of which are not observable. Nevertheless, we expect the latent variables extracted by VAE to provide the computational basis for further understanding the origins of resting state activity. We hypothesize that the truly disentangled physiological origins, whether observable or not, are individually describable as the latent variables up to linear and sparse projection. This hypothesis awaits confirmation by future studies.

In the latent space, functional connectivity between latent variables describes the geometry of the latent representation of rsfMRI activity. This is a new perspective different from the functional connectivity among observable voxels, regions or networks (Biswal et al., 1995; Yeo et al., 2011). If the VAE model has fully disentangled the sources in a population level, functional connectivity should be near zero between different latent variables and thus reflect a spherical geometry. In other words, the model sets a nearly null population-level baseline, against which individual variation stands out. The latent-space functional connectivity given data from a single subject becomes a unique feature of that subject. Supporting this notion, the use of functional connectivity in the latent space allows for a significantly improved accuracy, robustness, and efficiency in individual identification, compared to the use of functional connectivity among cortical parcels (Amico and Goñi, 2018; Byrge and Kennedy, 2019; Finn et al., 2015; Mejia et al., 2018; Venkatesh et al., 2020).

Note that our main purpose is not to push for a higher identification accuracy but to understand the distribution and geometry of data representations in the feature space. Therefore, we opt for minimal preprocessing and the simplest strategy for individual identification. There is room for methodological development to further improve the identification accuracy or to extend it for many other tasks, including classification of the gender or disease states, prediction of behavioral and cognitive performances, to name a few examples. We expect that such applications would be fruitful and potentially impactful to cognitive sciences and clinical applications.

---

**Supplementary Figure 1.**
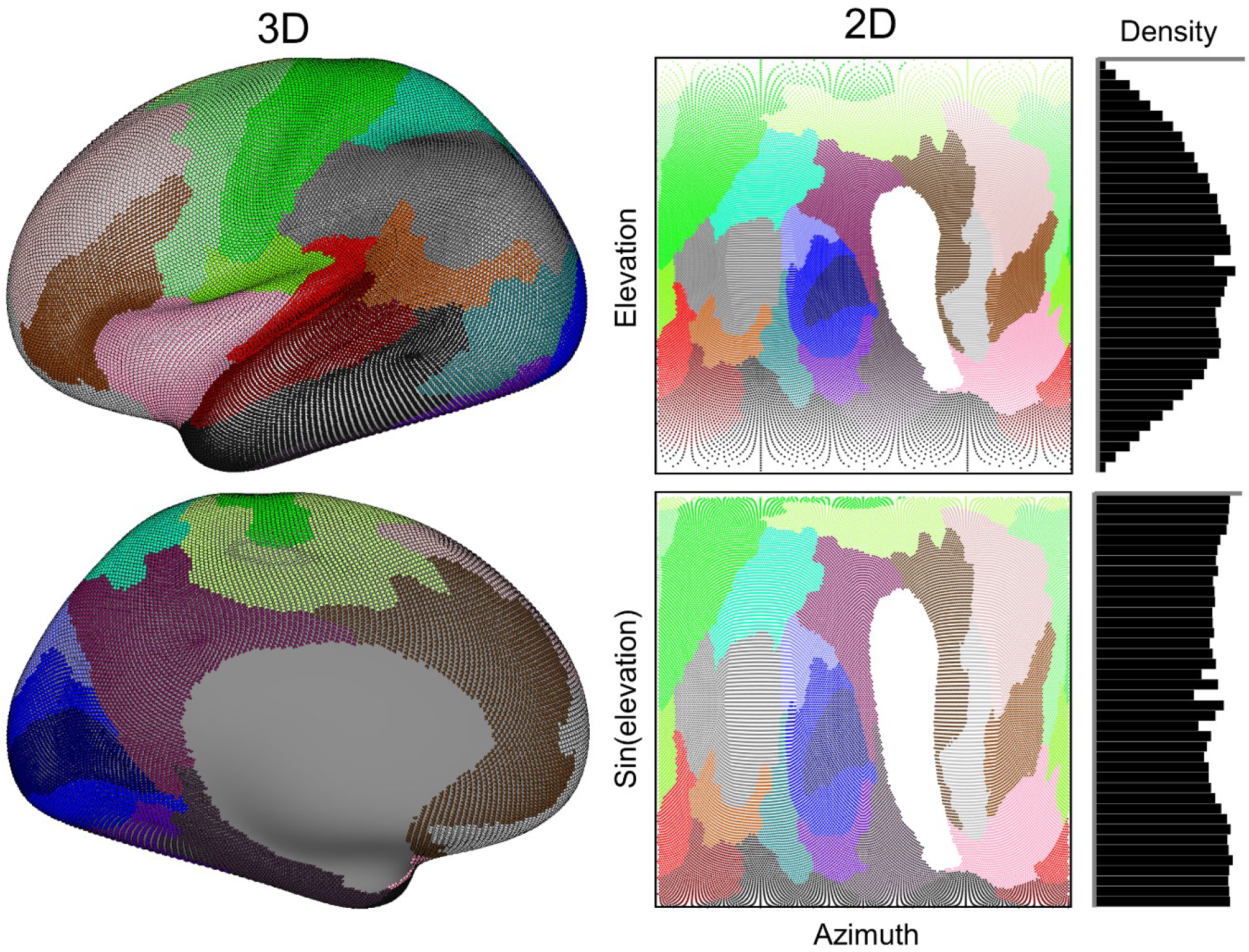
Grayordinates (# = 29,696) in left hemisphere (left panel) is projected into 2D space (middle panel) based on their azimuth- and elevation levels. Each color stands for different brain atlas based on the literature (Glasser et al., 2016). The imbalance in data density along the varying elevation level (upper in right panel) is alleviated by further applying elevation to Sine function (bottom in right panel).

**Supplementary Figure 2.**
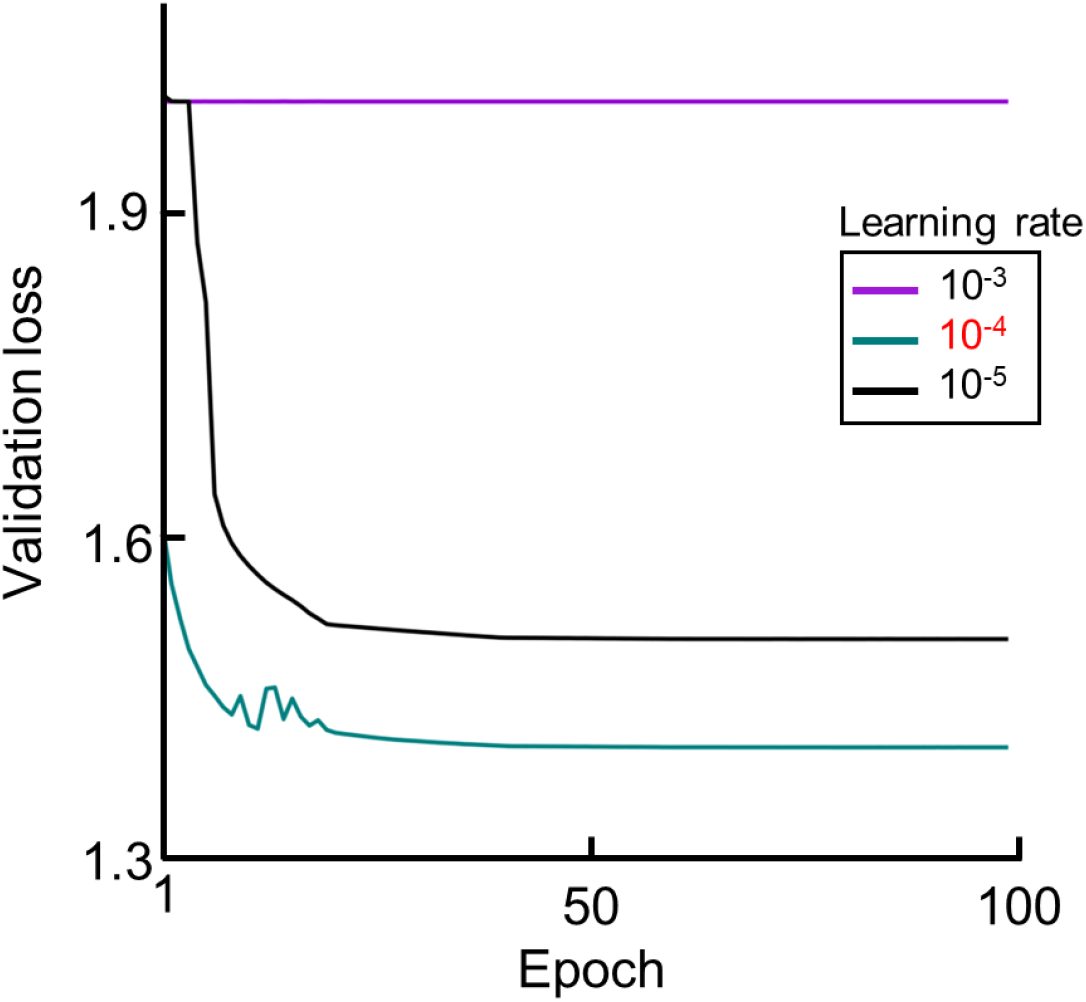
Validation curve of VAE at varying learning rate.

**Supplementary Figure 3.**
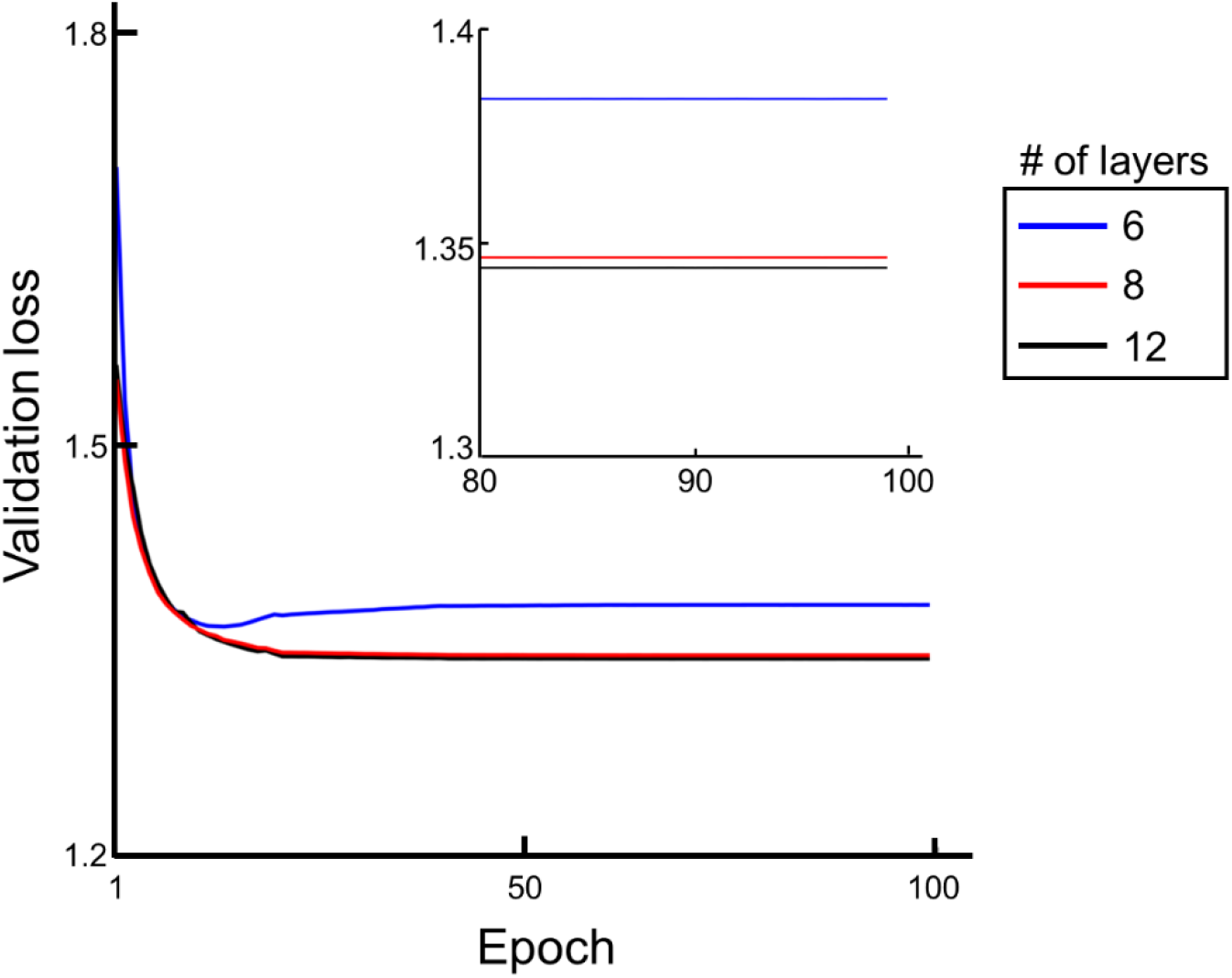
Validation curve of VAE at different layers.

## Notes

### Competing Interest Statement

The authors have declared no competing interest.

